# ER-stress signaling and Alzheimer’s proteins adjust the quality of human protein synthesis

**DOI:** 10.64898/2026.02.23.707382

**Authors:** Zhouli Cao, Max Hartmann, Maximilian Wagner, Amy L. Schug, Reinhild Roesler, Sebastian Wiese, Qingwen Yang, Franz Oswald, Karin Scharffetter-Kochanek, Sebastian Iben

**Author notes:** These authors contributed equally to this work (Zhouli Cao, Max Hartmann).

## Abstract

Proteostasis is the balance of protein synthesis, protein maintenance and protein degradation. Proteostasis is disturbed in neurodegenerative disorders like Alzheimer’s disease (AD) of the aging human body. Protein synthesis by the ribosome is the most error-prone process in gene expression. If and how the error-rate of protein synthesis is regulated during human aging and contributes to AD is unknown. Here we show that ribosomal error-rate is adapted in cellular models of human aging, but not in mouse aging. This adaptation involves ER-stress signaling and the Alzheimer’s disease-related proteins amyloid-beta precursor protein and presenilin 1. Our results suggest that ribosomal error-rate is a relevant parameter in human aging and disease.

## Introduction

Protein homeostasis (proteostasis) is essential for the proper functioning of cells, tissues, and organs in humans. Errors during gene expression could challenge proteostasis. During DNA replication, the error-rate is 10^-8^, the error-rate of gene transcription is 10^-6^, and protein synthesis by the ribosome produces 10^-4^ to 10^-3^ errors, creating misfolded proteins prone to aggregation ^1^. Protein synthesis errors are relevant to longevity - there is a striking linear correlation between ribosomal error-rates and lifespan in rodents ^2^. Moreover, raising the error-rate of the ribosome in mice leads to a premature aging phenotype, signs of Alzheimer’s disease (AD), and early death ^3,4^. Introducing a ribosomal mutation that reduces ribosomal error-rates extends the lifespan of *S*.*pombe, C*.*elegans* and *D*.*melanogaster* ^5^. Studies on ribosomal error-rates in human aging are sparse and mostly date back to the last century ^6,7^. Currently, it is unresolved whether translational errors by the ribosome contribute to the loss of proteostasis observed in AD and other neurodegenerative disorders ^8^. In an attempt to fill this gap, we asked whether and how the error-rate of protein synthesis is regulated during human aging, and whether Alzheimer’s proteins participate in this process.

## Results

### Reduced ribosomal error-rate and signs of ER stress in aging human fibroblasts

Translational fidelity, the accuracy of protein synthesis by the ribosome, is assessed with luciferase reporter assays. For this purpose, we introduced various inactivating mutations into the Nanoluciferase plasmid (near-cognate mutation and stop codons), and are now able to monitor errors in translation by the reactivation of Nanoluciferase activity ^9^. Our study utilized dermal fibroblasts from both young and old human donors. From the same donors, we also studied fibroblasts aged *in vitro* by serial passage. To explore whether the ribosomal error rate increases with human aging, as recently reported in mice ^10^, we examined our cellular models (Figure 1A). Interestingly, we observed a decreased error-rate in protein synthesis in both aging models, suggesting an increase in translational fidelity with human aging. This result was not due to differences in reporter expression efficiency, as revealed by qPCR (Supplemental Figure 1A). We further assessed protein synthesis using two methods: metabolic labeling with ^35^S-methionine after brief methionine starvation and measuring incorporation rates of o-propargyl-puromycin, analyzed via flow cytometry. These experiments demonstrated a significant decline in overall protein synthesis with age in both models (Figure 1B), although the levels of selected translation initiation factors remained unchanged (Supplemental Figure 1B). Overall protein synthesis is regulated by the phosphorylation state of the translation-initiation factor eIF2alpha. Increased phosphorylation of eIF2α suppresses translation initiation, and cells from older donors or late passages show higher levels of eIF2α phosphorylation as compared to those from young donors or early passages (Figure 1C). This increase was not associated with elevated translation from internal ribosomal entry sites (Supplemental Figure 1C). Phosphorylation of eIF2α indicates ER-stress or activation of the integrated stress response (ISR), as shown by the increase in ER-stress markers like Chop and ATF4 in fibroblasts from older donors or late passages (Figure 1D). ER-stress, caused by the accumulation of misfolded proteins, is sensed in the ER by the chaperone Binding Immunoglobulin Protein/Glucose Regulated Protein 78 kDa (BiP/GRP78), which binds and inactivates ER-stress effectors such as ATF6, IRE1α, and PERK. Misfolded proteins sequester GRP78 from these effectors, triggering the UPR. We observed reduced BiP/GRP78 protein levels in fibroblasts from older donors (Figure 1E), but not in late-passage fibroblasts. qPCR analysis showed decreased BiP/GRP78 mRNA in cells from older donors but increased levels in late-passage cells, suggesting that the amount of BiP/GRP78 alone does not control ER-stress response (eIF2α phosphorylation) in either aging model. The expression level of PERK mRNA was elevated in both models, as revealed by qPCR (Supplemental Figure 1D). Next, we examined whether there were more misfolded, aggregation-prone proteins in cell lysates from older donors compared to younger donors, which could explain the higher p-eIF2α. After applying heat shock to induce protein aggregation, the pelleted proteins were quantified. As shown in Figure 2F, there was even a trend toward fewer aggregation-prone proteins in lysates from both old donor-derived cells and high-passage cells. It is therefore tempting to speculate that the lower error rate of ribosomal translation shown in Figure 1A could affect the amount of aggregating proteins in old and late-passage cells. These findings suggest that the ribosomal error-rate decreases with human aging, along with reduced overall protein synthesis, signs of increased ER-stress, and a stable proteome.

**Figure 1.**
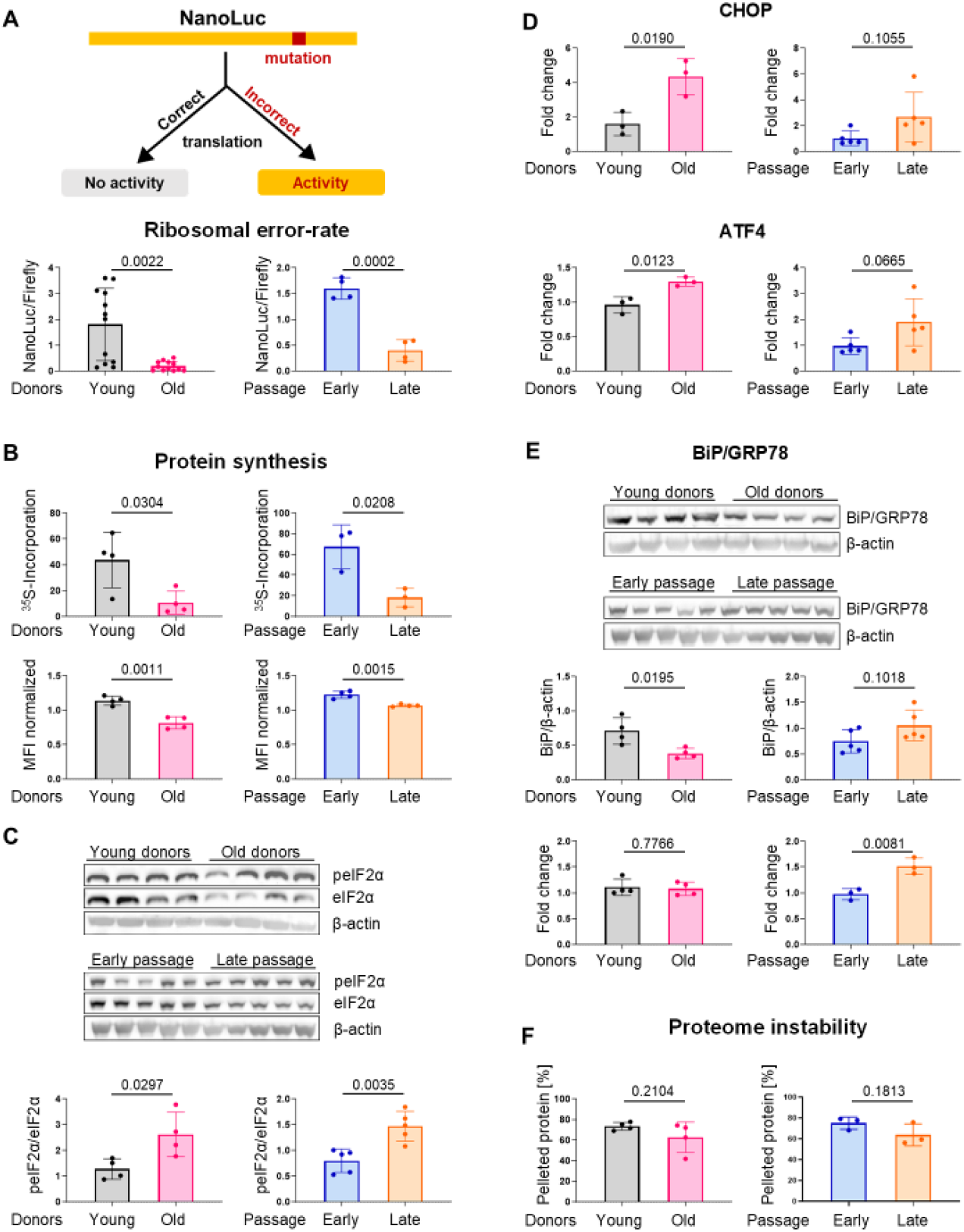
Ribosomal error-rate changes with aging. Figure 1 **A**. Principle of the translational fidelity assay. The ribosomal error-rate of primary skin fibroblasts from young and old donors (n = 4 in 3 independent experiments) (left), the error-rates of early and late passage primary fibroblasts (n = 4 independent experiments) (right). **B**. Protein synthesis assay by ^35^S-methionine incorporation (upper part), and by flow cytometry-based OPP-labeling (lower part). Each data point corresponds to an individual donor (n = 4) or an independent experiment (n = 3). **C**. Western blot analysis of eIF2α, phosphorylation with corresponding quantification. Each band corresponds to an individual donor (n = 4) or an independently cultured (n = 5) early- or late-passage fibroblast. **D** RT-qPCR analysis of UPR transcription factors CHOP and ATF4. Each data point corresponds to an individual donor (n = 3) or an independently cultured (n = 5) early- or late-passage fibroblast. **E**. Western blot and qPCR analysis of BiP/GRP78 from fibroblasts of young and old donors (n = 4), and early and late passages (n = 5, independently cultured fibroblasts). **F**. Proteome stability after heat treatment. Each data point corresponds to an individual donor (n = 4) or an independent experiment (n = 3) with early- or late-passage fibroblasts.

**Figure 2.**
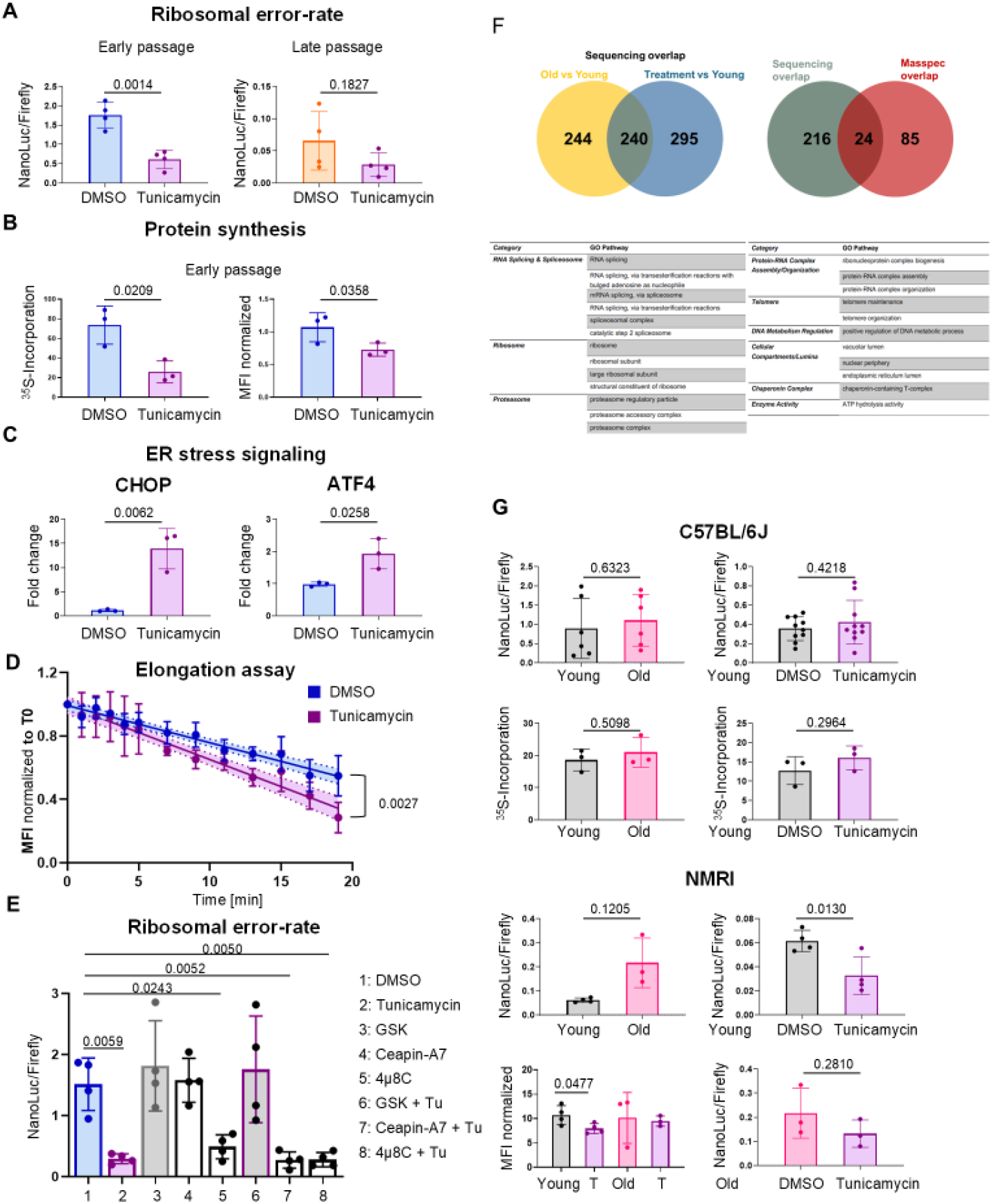
Ribosomal error-rate is affected by ER-stress signaling. A. Ribosomal error rate in early passage (left) and late passage (right) fibroblasts treated with tunicamycin (1 µg/ml) ( n = 4 independent experiments). B Protein synthesis analysis: incorporation of 35S-methionine (left) and OPP-labeling (right) in early passage fibroblasts treated with tunicamycin (1 µg/ml) for 24 hours (n = 3 independent experiments). C. Induction of CHOP and ATF4 expression by qPCR after tunicamycin treatment (n = 3 independently cultured cells). D. Elongation assay-kinetics of OPP-incorporation after blocking re-initiation by harringtonine (n = 3 independent experiments). E. Translational fidelity experiment of early passage fibroblasts with tunicamycin (1 µg/ml), PERK inhibitor GSK2606414 (1 µM), ATF6 inhibitor CeapinA7 (20 µM), IRE1α inhibitor 4µ8C (30 µM), and combined treatments (n = 4 independent experiments). F. Venn diagrams depicting the overlap of enriched GOs in old cells and tunicamycin-treated young cells (Nanopore sequencing left). The resulting 240 GOs were then compared with the mass spec overlap (right). The 24 resulting GO terms are listed. G. Translational fidelity assay with n = 6 (left), and n = 4 in 2 independent experiments (right), and protein synthesis (n = 3) in fibroblasts from young (12 weeks) and old (>100 weeks) C57BL/6J and NMRI mice.

### ER-stress regulates ribosomal error-rate

ER-stress signaling and the UPR have evolved to balance proteostasis and to reduce the amount of *de novo*-synthesized proteins when an imbalance of misfolded proteins in the ER is detected. To test whether provoking ER-stress impacts translational fidelity, we added tunicamycin to fibroblasts from early and late passages. Tunicamycin triggers ER-stress by inhibiting N-linked glycosylation, the initial step in glycoprotein synthesis. As shown in Figure 2A, the error-rate of ribosomes decreases after tunicamycin treatment in early passage cells, but not significantly in late passage cells, supporting the idea that late passage cells have already adjusted translational fidelity to minimize errors. Inducing ER-stress with thapsigargin produced a similar response (Supplemental Figure 2A) that was not affected by different reporter expression (Supplemental Figure 2B). Additional ER-stress, presumably transmitted via the same pathway, does not significantly further improve the error rate. Tunicamycin treatment is associated with a reduction in total protein synthesis (Figure 2B) and activation of UPR signaling (Figure 2C). EIF2α, when phosphorylated at serine 51, acts as a translational repressor by preventing the formation of the 43S translation initiation complex. However, speculating that elongation dynamics, as shown before^11^, may impact error-rate, we examined elongation using harringtonine and o-propargyl-puromycin incorporation. Harringtonine blocks re-initiation of translation, and the decline in the o-propargyl-puromycin signal reflects the elongation rate. Figure 2D shows that the signal in tunicamycin-treated cells (normalized to T0) drops more rapidly than in control cells, indicating that elongation speed even increases with higher ER stress. This result may suggest that reduced overall translation might lead to a better allocation of correct tRNAs for protein synthesis, thereby explaining the reduced errors. To identify the UPR effectors that enhance translation accuracy after ER-stress, we used a pharmacological approach to inhibit the functions of ATF6, IRE1α, and PERK. As shown in Figure 2E, tunicamycin treatment significantly lowers the error-rate in cells from young donors. The inhibitors GSK2506414 (PERK) and Ceapin A7 (ATF6) do not affect the basal error-rate, whereas the IRE1α inhibitor impacts translational fidelity. However, the PERK inhibitor GSK completely prevents the tunicamycin-induced decrease in error-rate, indicating that PERK activity helps reduce translation errors.

Next, we asked whether ER-stress induces transcriptomic and proteomic changes that are typical for old cells. We performed Nanopore sequencing and mass spectrometry analyses on fibroblasts from old versus young donors, then compared the results with those of young donors treated with tunicamycin (1 µg/ml for 24 hours). PCA analysis shows distinct separation between the groups in the transcriptomic data, and in the mass spectrometry analysis, the treated group clusters between the young and old donors (Figure S2C left, right). 50% of the transcriptomic GO entries enriched in old versus young cells were also enriched in tunicamycin-treated cells (Figure 2F). These GO entries were then compared to similar processed proteomic data, identifying 24 shared GO entries. Pathways such as ribosomes and RNA splicing suggest that cellular gene expression shifts toward posttranscriptional programs, indicating an aged phenotype following ER-stress.

Next, we investigated dermal fibroblasts from different mouse strains and did not detect a reduction in translational error-rate or protein synthesis with age, consistent with a recent publication ^10^. We observed a non-significant increase in the error-rate and a resistance to tunicamycin treatment in the C57BL6J strain (Figure 2G, Supplemental Figures 2 DEF). The reduction of ribosomal error-rate with age might be a feature of long-lived species as humans.

### Alzheimer proteins participate in the regulation of ribosomal error-rate

ER-stress and a loss of proteostasis are hallmarks of aging-associated neurodegenerative diseases ^12^. Therefore, we next asked whether the adjustment of ribosomal error-rate with aging involves the proteins Amyloid-beta precursor protein (APP) and the subunit of gamma-secretase that cleaves APP, presenilin1 (PSEN1). Mutations in both proteins can provoke early-onset Alzheimer’s disease (AD). We decided to knock down these proteins in immortalized fibroblasts by shRNA. The partial knockdown of both proteins in fibroblasts was successful, as shown by the Western blots in Figure 3A. Interestingly, this knockdown affected ER-stress signaling; APP knockdown increased eIF2α phosphorylation (Supplemental Figure 3A, right panel). The analysis of translational fidelity showed that knocking down APP decreased the error-rate of ribosomes and overall protein synthesis. This led to reduced proteome instability, as validated by the amount of aggregating proteins after heat shock. In contrast, knocking down PSEN1 decreased the phosphorylation of eIF2α and resulted in a “youthful” protein synthesis with a high error-rate, increased total protein synthesis, and an increased proteome instability. Employing HEK cells as a genetically easy-to-manipulate cell line, we confirmed our results after CRISPR/Cas9-mediated knockout (KO) of APP and PSEN1 (Supplemental Figures 3B/C). Using the HEK overexpression system, we transfected both wild-type (wt) and mutant APP into HEK cells and assessed translational fidelity. The “Swedish” mutation of APP, a double mutation causing amino acid substitutions at two neighboring residues (K670M and N671L), is associated with early-onset Alzheimer’s disease in humans ^13^. Overexpression of wt APP protein significantly increased the error-rate, while the mutant form of APP did not impact translational fidelity (Figure 3B and Supplemental Figure 3D). This might highlight that the “Swedish” mutation results in the loss of the important APP function, which regulates protein synthesis quality. Next, we performed a partial knockdown of APP and PSEN1 with shRNA in HEK cells and observed a comparable effect on ribosomal error-rate as in fibroblasts (Figure 3C). Notably, the regulator of ER-stress signaling, BiP, was strongly induced by PSEN1 knockdown (Figure 3C and Supplemental Figure 3E). BiP is known to stabilize APP ^14^ and might, in turn, be stabilized by more APP in the absence of APP-cleavage by PSEN1 knockdown. To further explore how BiP affects translational fidelity, we used shRNA to reduce BiP levels and assessed the impact of silenced BiP on ribosomal error-rate. In fact, BiP knockdown results in a noticeable decrease in the ribosomal error-rate (Figure 3D). The cell may detect BiP knockdown as ER-stress, potentially activating the UPR. Overall, these results reveal a new aspect of human APP and PSEN1 proteins that strongly impact proteome stability by adjusting translation error-rate and overall protein synthesis.

**Figure 3.**
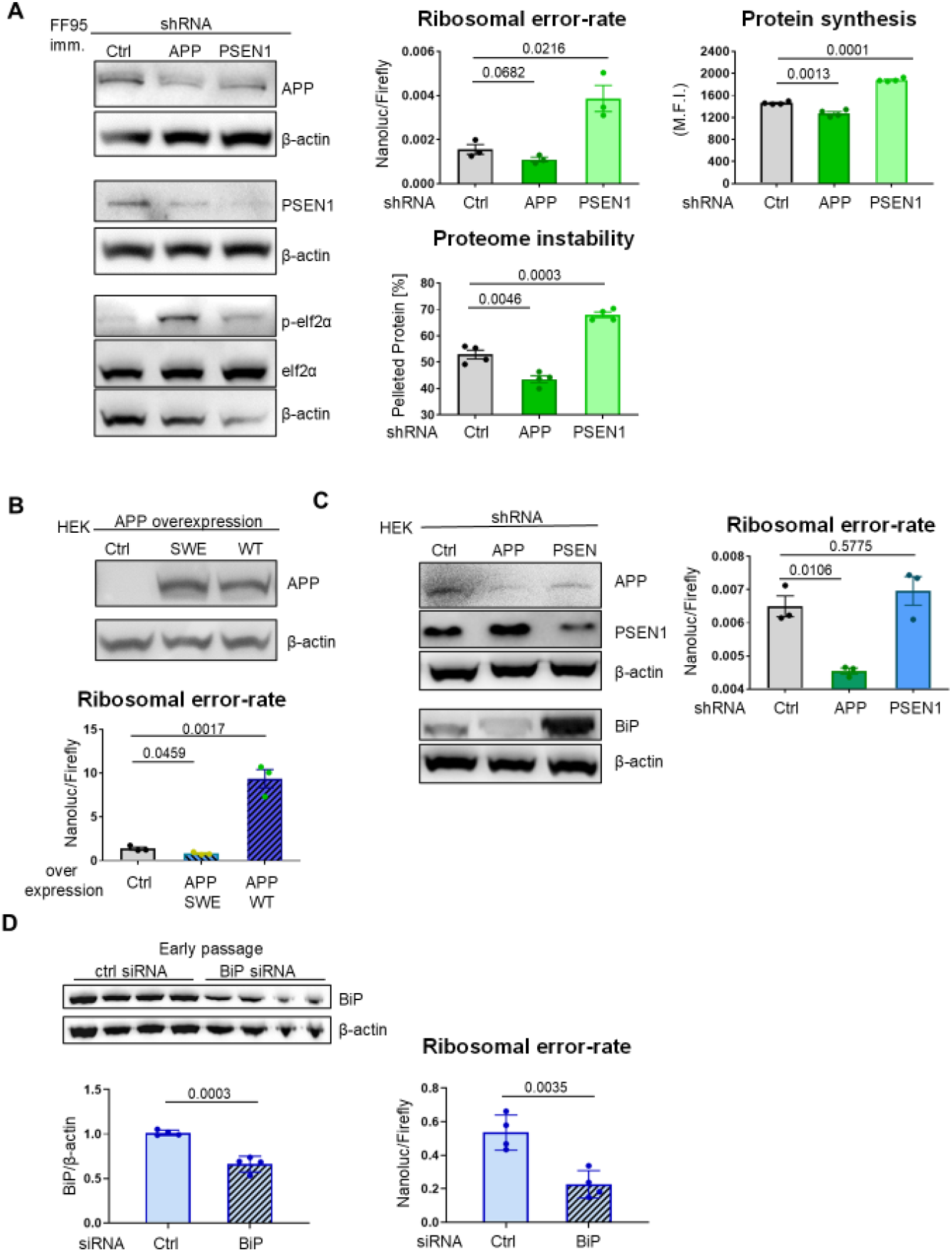
APP and PSEN1 are involved in the regulation of translational fidelity. **A**. Downregulation of APP/PSEN1 by shRNA in immortalized fibroblasts was demonstrated by Western blot and analyzed for translational fidelity (n = 3), protein synthesis (n = 4), and proteome instability (n = 5). Each data point represents an independent experiment. **B**. Overexpression of APP-SWE (Swedish mutation) and WT-APP, and ribosomal errors in HEK293T cells. Each data point represents an independent experiment (n = 3). **C**. Downregulation of APP/PSEN1 by shRNA in HEK293T cells and its impact on the ribosomal error rate. Each data point represents an independent experiment (n =3). **D**. Downregulation of BiP in early passage fibroblasts (Western blot) affects ribosomal error-rate (translational fidelity assay). Each data point represents independently cultured fibroblasts (n = 4).

### Pharmacological modulation of ribosomal error-rate

Next, we aimed to test interventions that affect ribosomal error rates and initially examined an inhibitor of PERK (GSK2606414). The addition of 1μM of GSK inhibitor unexpectedly but consistently lowered the ribosomal error rate in control cells across two different cell lines. However, APP knockdown in cells that previously decreased the error rate, could be reversed by PERK inhibition, as shown in fibroblasts (Figure 4A) and HEK cells (Figure 4B). These results indicate that the effect of APP reduction on the ribosomal error-rate is mediated through PERK. To further verify that the effect of APP manipulation on the error-rate is specific, we used CRISPR/Cas9 to knock down Notch, another gamma-secretase target. Notch knockdown with two different guide RNAs did not impact ribosomal accuracy (Figure 4C). We then tested whether gamma-secretase inhibitors also affect error-rates. After confirming the effects of these inhibitors on Notch activity (Supplemental Figure 3F), we treated human fibroblasts with the inhibitors. Two of the three inhibitors significantly increased the error-rate, mimicking PSEN1 knockdown (Figure 3D). This effect was entirely dependent on APP presence, as knocking down APP prevented the effect of gamma-secretase inhibitors, indicating that APP is essential for this regulation. These results could not be replicated in mouse cells, APP knockdown or -secretase inhibition had no effect on ribosomal error-rate (Figure 4E), suggesting that APP and PSEN1 have gained additional functions in human cells. Moreover, a low dose of the translational inhibitor cycloheximide extends the lifespan of *C. elegans* ^15^, leading us to hypothesize that cycloheximide might also influence ribosomal error-rates. Of note, nanomolar concentrations of cycloheximide significantly decreased the error rate without impairing protein synthesis (Figure 4F). Overall, our findings suggest that in human cells, APP directly impacts ER-stress signaling and helps maintain translational accuracy.

**Figure 4.**
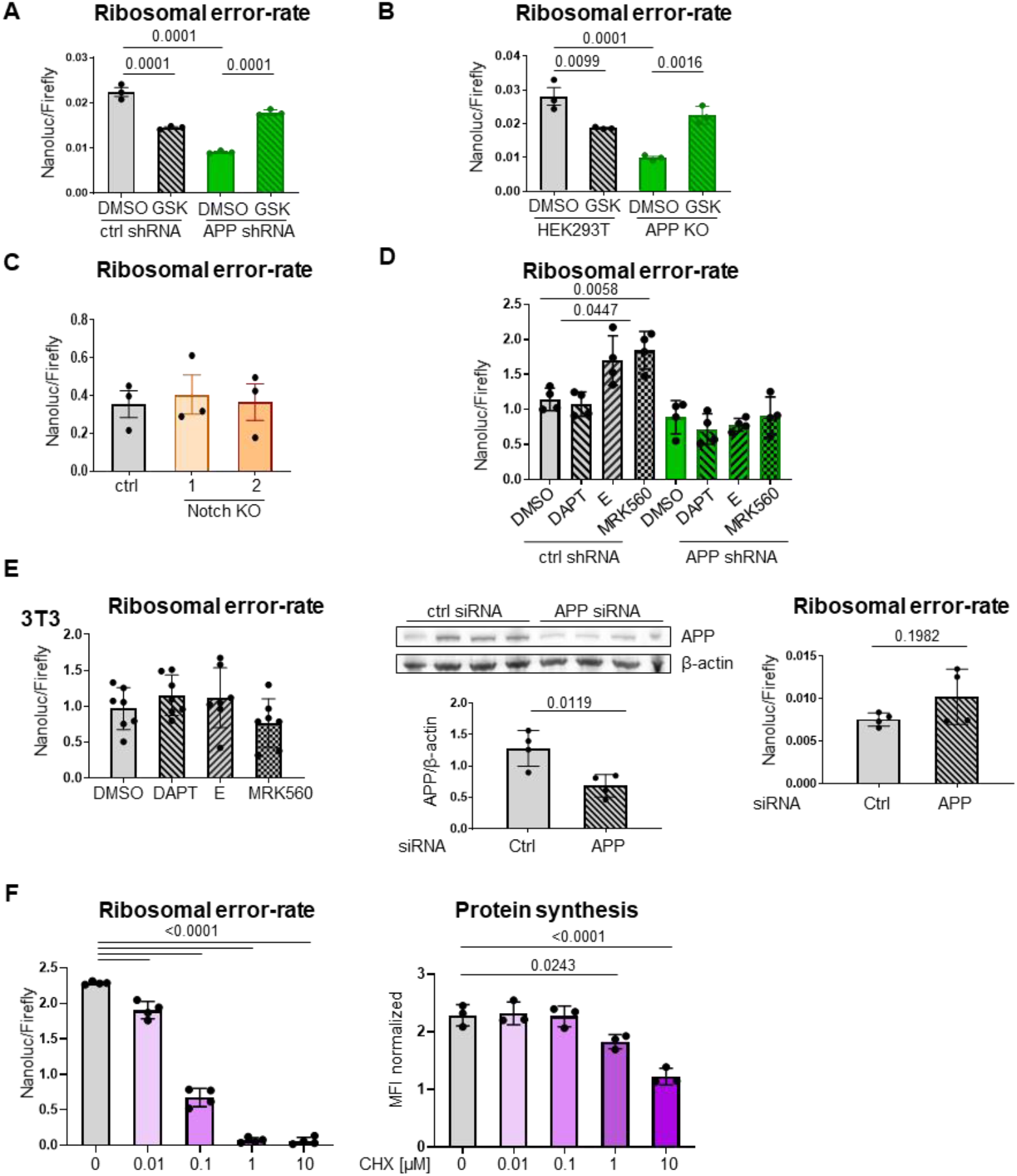
Translational fidelity experiments reveal regulation-dependence on human APP. **A**. Effect of PERK inhibition by GSK2606414 (1 µM) in both shRNA APP knockdown and control shRNA fibroblasts (immortalized) on the ribosomal error-rate. N = 3, independent experiments. **B**. Same experiment as Panel A, but in an APP-KO HEK293T cell line. N = 3, independent experiments. **C**. General knockdown of Notch does not influence the ribosomal error-rate. N = 3, independent experiments. **D**. The effect of PSEN1 inhibitors (DAPT 10µM, Compound E 5µM, MRK560 1µM) on the ribosomal error-rate in immortalized fibroblasts with or without APP downregulation. N = 4, independent experiments. **E**. Ribosomal error-rate in fibroblasts from 3T3 mice treated with PSEN1 inhibitors. N =7, independent experiments. APP downregulation with siRNA and measurement of ribosomal error rate in 3T3 cells. N = 4, independently cultured cells. **F**. Low-dose cycloheximide (CHX) treatment affects ribosomal error-rate (left). N = 4, individual donors. Impact of low-dose cycloheximide (CHX) treatment on protein synthesis (right). N = 3, individual donors.

## Discussion

Ribosomal errors during translation have long been discussed as an aging parameter. An aging hypothesis based on ribosomal errors, called the “error catastrophe theory of aging,” was proposed by Leslie Orgel in the 1960s ^16^. It became increasingly clear that translational fidelity influences health and lifespan in animals ^2,3,5^. Goldstein et al. already noted a decreased error rate in *in vitro*-aged human fibroblasts ^17^. In fact, ribosomal error-rates in long-living humans, but not in short-living mice, are regulated. This may be an adjustment to the declining cellular and organismal compensation capacity ^18^ of the aging human body ^8^, thereby contributing to healthy aging. A constant or rising error-rate in the protein synthesis machinery with age could overwhelm the declining maintenance and degradation capacities of aging cells and tissues, resulting in the accumulation of misfolded proteins. Our data show that the youthful state, with unrestrained cellular and organismal compensation capacity, is characterized by a high error-rate but high protein synthesis, whereas older cells make fewer errors along with reduced protein synthesis. Thus, human aging can be viewed as a delicate balance between the need for *de novo* protein synthesis to support memory and cognition ^19^ and the costs of high protein synthesis, such as errors, misfolded proteins, and protein aggregation.

Comparative transcriptomic studies of 103 mammalian species identified pathways related to translational fidelity correlating with longevity ^20^. Translational fidelity varies between mice and humans ^21^, and we here confirmed that the error-rate increases with mouse aging. Currently, it is unclear whether the mechanism of a reduced error-rate with aging is a private or general mechanism of long-lived species.

Our results show that the Amyloid-beta precursor protein APP influences the quantity and quality of protein synthesis in human cells, but not in mice. Likely through its interaction with BiP/GRP78 ^14^, which acts as the sensor and effector for misfolded proteins. It has previously been reported that ER-stress induces APP degradation ^22^, supporting our hypothesis that reduced APP levels result in less BiP stabilization and increased PERK signaling. This leads to eIF2α phosphorylation, lower overall translation, and fewer errors, thereby decreasing the ER’s load of misfolded proteins. Consistent with this idea, -secretase cleavage of APP appears to reduce ribosomal errors and total translation. Pharmacological inhibition of □-secretase would then cause APP accumulation, BiP stabilization, and decreased PERK activity. This reflects the “youthful” translation state, characterized by high total translation, elevated error-rates, and proteome instability. It is plausible to speculate that this condition is harmful in diseases marked by proteome instability, such as AD. Here, we identified a novel ER-stress signaling cascade involving human APP, affecting ribosomal accuracy and proteome stability, which may impact our understanding of Alzheimer’s disease.

## Acknowledgements

We thank Meinhard Wlaschek for stimulating discussions. Old donor cells were obtained from Hertie-Stiftung Tübingen. This work was funded by the German Research Foundation (DFG) program CRC Aging at Interfaces (1506 A05, A06, C02). Z.C. and Q.Y. are supported by the China Scholarship Council (CSC File No. 202206090008/202206090009)

## Authors contributions

Z.C.and M.H. performed most experiments and wrote a draft; M.W, A.S., R.R. Q.Y- and S.W. conducted specific experiments and analyses, F.O. and K:S-K. supervised the works, S.I. initiated, supervised and finalized the study and the manuscript.

## Supplementary materials

### Material and Methods

#### Cell culture

Human dermal fibroblasts, HEK293T, and 3T3 cells were cultured at 37°C with 5% CO_2_ in DMEM (Gibco, 41965-039) supplemented with 10% FBS (Sigma-Aldrich, S0615), 1% Penicillin-Streptomycin (PAN-Biotech, P06-07100), and 1% Glutamine (PAN-Biotech, P06-80100). Primary mouse fibroblasts were cultured under the same conditions but at 37°C, 5% CO_2_, and 3% O_2_. Fibroblasts from four different young donors were obtained from the foreskin tissue of donors under 1 year old. Fibroblasts from older donors ranged in age from 74 to 84 years. Early passage fibroblasts were maintained below CPD25, while late passage fibroblasts were cultured until reaching senescence (> CPD60). Isolated mouse fibroblasts were obtained from the ears of young (12 weeks) or old (>100 weeks) mice through digestion with Collagenase A.

#### CRISPR/Cas9-Mediated Gene Knockout via Electroporation

CRISPR/Cas9-mediated knockout of PSEN1 and APP was achieved by electroporating cells with 1 µg of Cas9/sgRNA plasmid and 1 µg of linear donor DNA carrying the EF1α-GFP-P2A-Puromycin selection cassette (Origene, KN409575 and KN416443). The DNA–cell mixture (10 µL, 1 × 10^7^ cells/mL) was loaded into a 10 µL Neon™ tip and electroporated using the optimized Neon™ parameters. Electroporated cells were immediately transferred to pre-warmed medium without antibiotics and cultured to recover. Cells were passaged at a 1:10 ratio starting 48 hours after electroporation and repeatedly diluted over 1–2 weeks to remove non-integrated donor DNA. Puromycin selection (1 µg/mL) was initiated after passage 3–5 and continued until unedited cells were eliminated. Gene disruption and donor integration were confirmed by Western blot or fluorescence-based analysis. Single-cell clones were obtained either by limiting dilution or colony picking and expanded for downstream experiments. sgRNAs used in this study

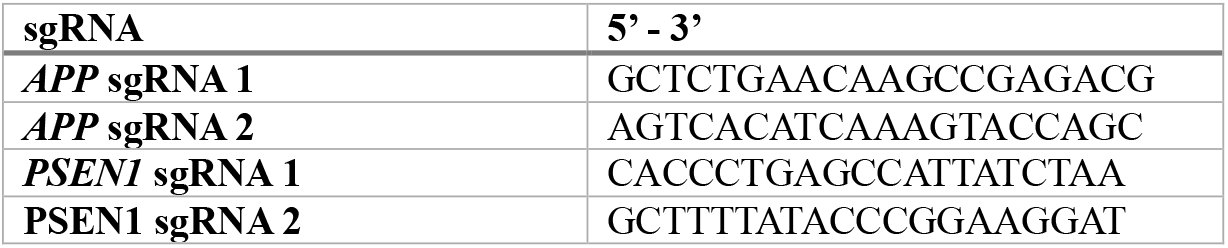

#### shRNA-Mediated Gene Knockdown via Electroporation

shRNA knockdown was achieved by electroporating 1 µg of shRNA plasmid (or non-targeting control) with 10 µL of cell suspension (1 × 10^7^ cells/mL) using the Neon™ Transfection System under the same electroporation conditions as used for CRISPR experiments. After electroporation, cells were transferred into antibiotic-free medium and allowed to recover for 18–24 hours. For transient knockdown, cells were collected 24–72 hours post-electroporation for downstream assays. For stable knockdown, puromycin selection (1 µg/mL) was initiated 48 hours after electroporation and maintained by changing the medium every 2–3 days until all control cells were eliminated. Knockdown efficiency was assessed at the mRNA level by RT-qPCR and at the protein level by Western blot before proceeding with functional assays.

#### Translational fidelity assay

Transfection with the NanoLuc and Firefly reporters was performed as described in Hartmann et al. 2025. The effects of inhibitors or ER-stress inducers on the ribosomal error-rate were examined by adding the respective drug to the cell culture medium. NanoLuc and Firefly activity were developed 24 hours post-transfection.

All compounds that were used in this study are listed in the following table.

| Name | Purpose | Company | Catalogue number |
| --- | --- | --- | --- |
| **4µ8C** | IRE1 inhibitor III | MedChemExpress | HY-19707 |
| **Ceapin-A7** | ATF6 inhibitor | MedChemExpress | HY-108434 |
| **Compound E** | γ-secretase inhibitor | Sigma-Aldrich | 565790 |
| **DAPT** | γ-secretase inhibitor | Sigma-Aldrich | D5942 |
| **GSK2606414** | PERK inhibitor | MedChemExpress | HY-18072 |
| **MRK560** | γ-secretase inhibitor | MedChemExpress | HY-14174 |
| **Thapsigargin** | ER stress inducer | MedChemExpress | HY-13433 |
| **Tunicamycin** | ER stress inducer | Cell Signaling | 12819S |

#### siRNA transfection

One day before the experiment, 1000000 cells were seeded into a 6 cm dish. Lipofectamine 3000 (Thermo Fisher, L3000008) was used according to the manufacturer’s protocol for transfection. 48 hours post-transfection, cells were collected for Western blot and translational fidelity assay. In this study, siRNAs against BiP (Santa Cruz, sc-29338), against mouse APP (sc-29678), and control siRNA (sc-37007) were used.

#### *In vitro* translation

For *in vitro* translation, 20 µl of cytosolic lysate (lysed with hypotonic buffer), 4 µl of 10x translation mix buffer (152 mM HEPES pH 7.5, 8 mM ATP dipotassium salt, 480 µM GTP disodium salt, 2 mM spermidine, 16 mM DTT, 80 mM creatine phosphate, 200 µg/ml creatine phosphokinase, 200 µg/ml calf liver tRNA, 100 µM amino acid mix), and 600 ng of mRNA were combined and then brought up to 40 µl with water. The mRNAs used in this study are synthesized from the linearized plasmids. The pCLneo-CMV-DualLuc-K529N, provided by Dr. Markus Schosserer, allows measurement of the ribosomal error rate. The CrPV-IRES (provided by Marianna Penzo) construct enables measuring both cap-dependent and cap-independent translation.

#### RNA extraction, cDNA synthesis, and RT-qPCR

RNA was extracted using the QIAGEN RNeasy® mini kit (cat. number 74106) following the manufacturer’s protocol. 1 µg of RNA was reverse transcribed into cDNA with M-MLV reverse transcriptase (Promega, M170B) according to the manufacturer’s instructions. For RT-qPCR, FastStart Universal SYBR Green Master (Roche, 04913850001) was used in a 30-cycle PCR program on the QuantStudio 5 real-time PCR system (Thermo Fisher).

All primers that were used for qPCR in this study are listed in the following table.

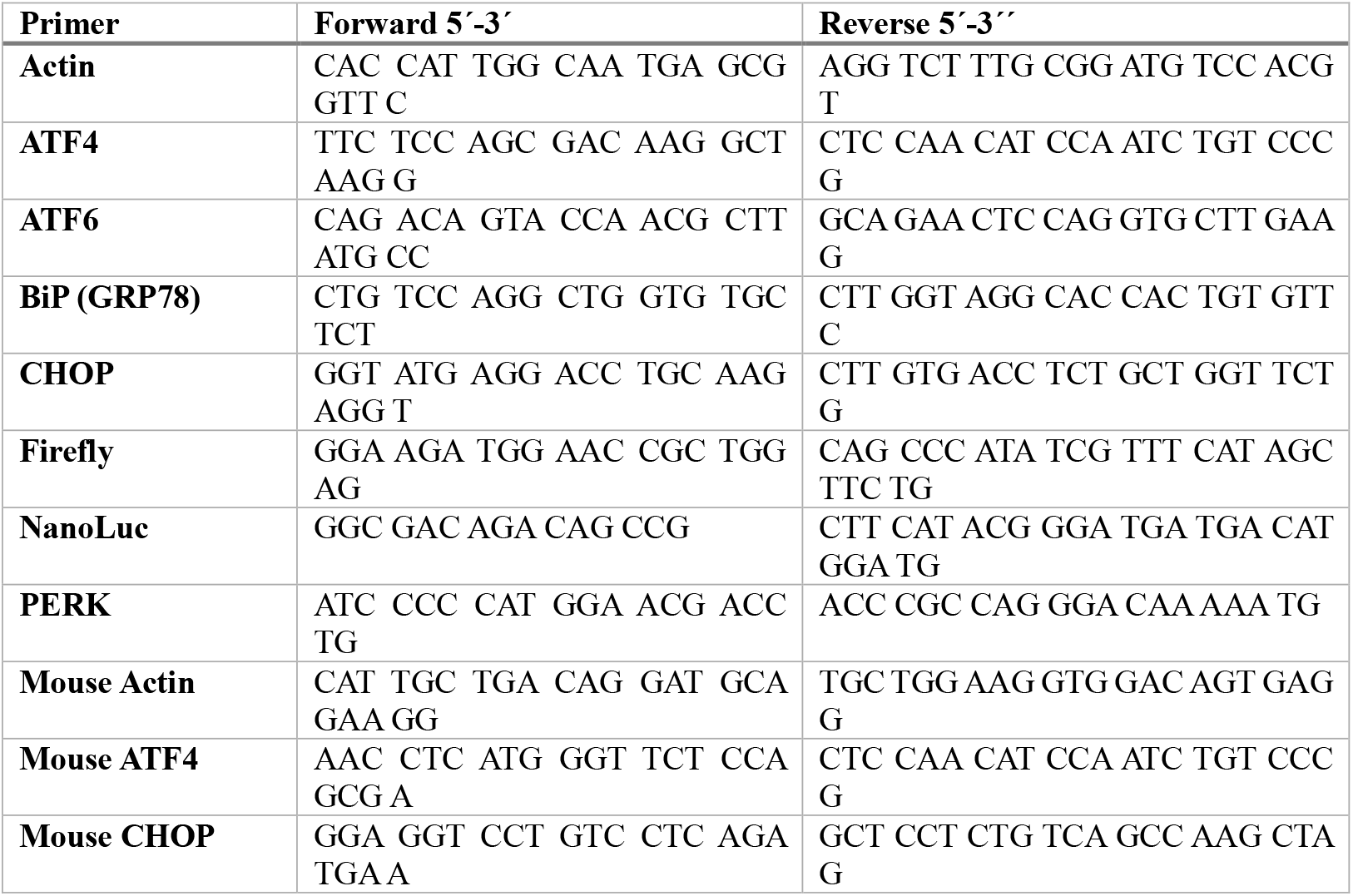

#### Western blot analysis

Collected cell pellets were lysed with a standard detergent-based lysis buffer supplemented with Halt™ protease & phosphatase inhibitor cocktail (100x) (Thermo Fisher Scientific, 78440). Bolt™ 4 to 12%, Bis-Tris, 1.0 mm, mini protein gels (Invitrogen, NW04122BOX) were used for SDS-PAGE. Proteins were transferred to an Amersham™ Protran™ 0.45 µm nitrocellulose membrane (MERCK, GE1060002) overnight at 4°C using a wet transfer at 30V. The membrane was blocked with 5% milk in TBS-T (Sigma-Aldrich C7078) or 5% BSA for 30 min, as recommended by the antibody manufacturer. Primary antibody incubation was performed overnight at 4°C in either 5% milk or 5% BSA. After washing and secondary antibody incubation, the Western blot was developed using the SuperSignalTM Western Blot Substrate (Thermo Fisher Scientific, A43840) with the Fusion Fx7 (Vilber) imaging system.

Each lane represents an individual donor or a replicate from an independent cell culture and lysate. Fiji ImageJ was used for Western blot quantification (densitometry analysis). Unphosphorylated proteins were detected on the same membrane as their phosphorylated counterparts. The exposure time for unphosphorylated protein detection was significantly shorter than that for the phosphorylated protein, ensuring that the phosphorylated proteins had little to no impact on the analysis of the unphosphorylated protein.

| Antibody | Company | Catalogue number |
| --- | --- | --- |
| **β-actin HRP** | Santa Cruz | sc-4778 |
| **BiP/GRP78** | Cell Signaling | 3177S |
| **p-eIF2α** | Cell Signaling | 9721S |
| **eIF2α** | Cell Signaling | 9722S |
| **PERK** | Santa Cruz | sc-377400 |
| **APP** | Thermo Fisher | PA5-17829 |
| **PSEN1** | Santa Cruz | sc-365450 |
| **eIF4G** | Cell Signaling | 2469T |
| **eIF4E** | Cell Signaling | 2067T |
| **p-4E-BP1** | Cell Signaling | 2855P |
| **4E-BP1** | Cell Signaling | 9644P |

#### O-Progargyl-Puromycin (OPP) labeling

The Protein Synthesis Assay Kit (Cayman Chemical, Cay601100-96) was used following the manufacturer’s instructions to measure protein synthesis. The incubation time of OPP was set to 20 minutes to prevent saturation of OPP incorporation. Analysis was performed using flow cytometry as described in the manufacturer’s protocol. The intensity of the OPP-labeled sample was normalized to the sample labeled with 5-FAM-azide-only to subtract background and possible intensity shifts due to differences in cell size across donors.

#### Elongation Assay

The elongation assay (SunRiSE assay) was adapted from Argüello et al., 2018. Instead of using puromycin and puromycin antibody, OPP was employed. Harringtonin (Santa Cruz, Sc-204771) was added at time zero, and OPP was added at the time points 0, 1, 2, 3, 4, 5, 6, 7, 9, 11, 13, 15, 17, and 19 minutes after harringtonine. Fibroblasts were cultured with tunicamycin (1 µg/ml) for 24 hours before the experiment. Flow cytometry was used to analyze protein synthesis.

#### Proteome instability assay

This assay was adapted from Treaster et al. 2014. Cytosolic extracts were ultracentrifuged for 1 hour at 100,000 g and 4°C to remove any aggregates. 100 µg of protein lysate was heated at 99°C for 15 minutes and then centrifuged for 5 minutes at 16900 g. The ratio of protein in the pellet compared to the supernatant (measured by Bradford) provides information about proteome instability.

#### Notch Cleavage Activity Assay

Notch cleavage activity was assessed using a firefly luciferase reporter system. The firefly reporter plasmid pGA98/6 was co-transfected transiently into cells with the ΔE Notch construct. After 4-6 hours, cells were treated with a γ-secretase inhibitor (Component E, DAPT, or MRK560) or DMSO (as a control). Cells were incubated for another 24–48 hours, followed by cell lysis and measurement of firefly luciferase activity. Luciferase activity was analyzed to quantify Notch cleavage, and inhibitor-treated samples were used to validate γ-secretase-dependent activation.

#### Nanopore sequencing

Oxford Nanopore transcriptome sequencing was performed as previously described, with minor modifications ^23,24^. For each of the 12 samples, 20 µg of high-quality total RNA (RIN ≥ 9) were used for library preparation with the Direct cDNA Sequencing Kit (#SQK-DCS109, Oxford Nanopore Technologies) in combination with barcodes and adapters from the Ligation Sequencing Kit (#SQK-LSK114). In brief, RNA was reverse-transcribed using the VNP poly(A) primer and SSP strand-switch primer with Maxima H Reverse Transcriptase (#EP0753, Thermo Fisher Scientific). The RNA template was then digested, and the second DNA strand was synthesized. After a clean-up step with AMPure XP beads (#A63881, Beckman Coulter), end-repair was performed using the NEBNext Ultra II End Repair/dA-Tailing Module (#E7546L, New England Biolabs).

Barcodes were ligated individually, and all samples were subsequently pooled into a single library. Native sequencing adapters were then ligated, followed by a final clean-up with Short Fragment Buffer (SFB). The completed library was eluted in 25 µl Elution Buffer (EB) and quantified using the Qubit High Sensitivity dsDNA Kit according to the manufacturer’s instructions. Two PromethION flow cells were primed following the standard Oxford Nanopore protocol, and 300 ng of the final library were loaded onto each flow cell. Sequencing was performed for 72 hours using default settings, with live basecalling disabled. Flow cells were washed twice (after 36 h and 50 h), and 300 ng of library were reloaded each time.

Raw data were basecalled post-run using Dorado v1.0.1 with the “sup” model and default parameters. Reads were aligned to the human reference genome (GRCh38), and transcript quantification was performed with bambu v2.2.0 ^24^. Samples were screened for outliers and potential batch effects, which were not detected. Given the high sequencing depth but limited sample size, gene expression estimates were treated as standardized continuous values and log-transformed before differential expression analysis. The limma framework was selected here to provide a robust statistical framework for normalized expression estimates and to ensure methodological consistency with downstream proteomic analyses, which were analyzed using the same approach. Multiple test corrections were applied to p-values using the Benjamini-Hochberg method to control for false discovery rates. Gene identifiers were standardized using an NCBI-based annotation module before downstream analysis. This module converts Ensembl or Entrez gene identifiers into standardized gene symbols while integrating curated gene description information. Only uniquely mapped genes were retained for subsequent analysis. Gene set enrichment analysis (GSEA) was performed using the ordinal gene list derived from the differential expression results. Genes were sorted by log2 fold change and analysed using the ‘gseGO’ function and ‘fgsea’ algorithm from the ‘clusterProfiler’ package. Enrichment analysis was performed on the biological processes, molecular functions, and cellular component categories of the Gene Ontology (GO). GO entries with a nominal p-value < 0.05 and a corrected p-value (false detection rate) < 0.25 were considered enriched. Enrichment results were interpreted cautiously and primarily used to identify shared functional trends. Pathways related to cell cycle regulation, chromatin organization, DNA replication and repair, ribosome biogenesis, and RNA processing were overlapped, while the result was not interpreted as direct evidence of tissue identity or developmental patterning.

### Mass spectrometric analyses

#### Sample preparation

50µl of DIGE-buffer (30 mM Tris-base, 7 M Urea, 2 M Thiourea, pH 8.5) was added to each sample pellet for cell lysis. Subsequently, 5ul of each sample was diluted in 30µl 50mM ammonium bicarbonate followed by reduction with 5 mM DTT (AppliChem, Darmstadt, Germany) for 20 min at RT, and alkylation with iodoacetamide (SigmaAldrich, St. Louis, USA) for 20 min at 37°C. Samples were digested overnight at 37°C using Trypsin at a 1:50 enzyme-to-protein ratio.

#### LC-MS/MS

15µl of each digest was measured using LTQ Orbitrap Elite system (Thermo Fisher Scientific) online coupled to an U3000 RSLCnano (Thermo Fisher Scientific) as described previously ^25^ with the following modifications: The column was initially 5 min equilibrated in 5% B (Solvents: A 0.1% FA; B 86% ACN, 0.1% FA). Followed by different elution steps, in which the percentage of B was first raised from 5 to 15% in 5 min, followed by an increase from 15 to 40% B in 105 min. The 20 most intense ions from the survey scan were picked for CID fragmentation. Singly charged ions were rejected, and the m/z of fragmented ions were excluded from fragmentation for 60s. MS2 spectra were acquired employing the LIT at rapid scan speeds. For each sample, two technical Replicates were measured.

#### Data analysis

Database search was performed using MaxQuant Ver. Ver. 2.4.13.0 (www.maxquant.org) ^26^. For peptide identification, the built-in Andromeda search engine ^27^ was employed to correlate MS/MS spectra with the UniProt human reference proteome set (www.uniprot.org). Carbamidomethylated cysteine was considered a fixed modification. False Discovery rates were set on both the peptide and protein levels to 0.01. Match between runs and dependent peptides search was enabled. Quantification was achieved based on intensities. The obtained data were processed by filtering out reverse hits. Graphs were generated using Origin Pro 2017G.

Mass spectrometry-based proteomics data were processed and analyzed using a workflow consistent with transcriptomics analysis to enable direct cross-platform comparisons. Raw mass spectrometry data were used for protein identification and quantification using standard proteomics workflows. Quantitative intensity values of protein levels were logarithmically transformed and normalized before statistical analysis. Differential protein abundance analysis was performed using the limma framework, which models continuous quantitative data and applies empirical Bayesian methods to stabilize variance estimates, particularly suitable for studies with limited sample sizes. Multiple test corrections for p-values were performed using the Benjamini-Hochberg method to control for false discovery rates. Samples were evaluated for potential outliers and batch effects, but none were detected. For integrative analysis, carefully curated annotation resources were used to map protein identifiers to gene symbols, ensuring compatibility with transcriptomics datasets. Only uniquely mapped proteins were retained for subsequent analysis. Functional interpretation of proteomic changes is performed in parallel with transcriptomic analysis, enabling direct comparison of pathway trends across different molecular levels.

#### Statistics

GraphPad Prism was used for statistical analysis of the experiments. In this study, each data point represents the average of technical replicates from an individual donor, an independent experiment, or an independent culture using the same cell line. First, a Shapiro-Wilk test confirmed that the data were normally distributed (Gaussian). For graphs comparing two groups, an F-test was used to assess variance. If the F-test showed non-significant variance, an unpaired, two-tailed Student’s t-test was performed. If the F-test showed significant variance, an unpaired, two-tailed Student’s t-test with Welch’s correction was performed. To compare more than two groups, a one-way ANOVA with multiple comparisons was performed. The p-values are always shown in the figure.

---

**figure S1.**
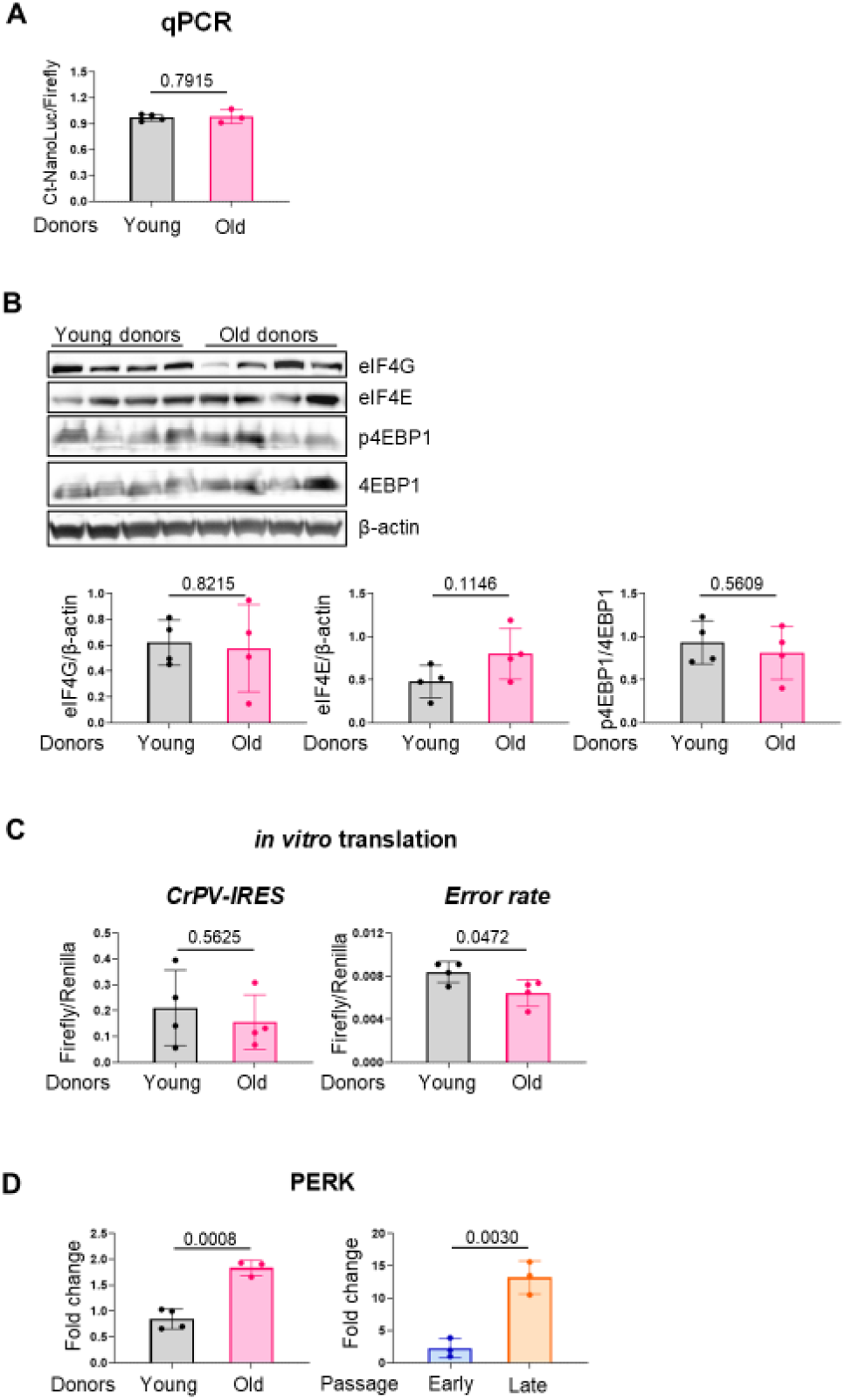
**A** Similar expression level of NanoLuc and Firefly luciferase after transfection, assessed by qPCR. Young n = 4, old n = 3, individual donors **B**. Western blot analysis of eukaryotic initiation factors in fibroblasts from young and old donors (n = 4). **C**. In vitro translation of CrPV-IRES and FireflyK529N/Renilla constructs in lysates from young and old donors (n = 4). **D**. RT-qPCR analysis of PERK in the aging models (n = 3).

**figure S2.**
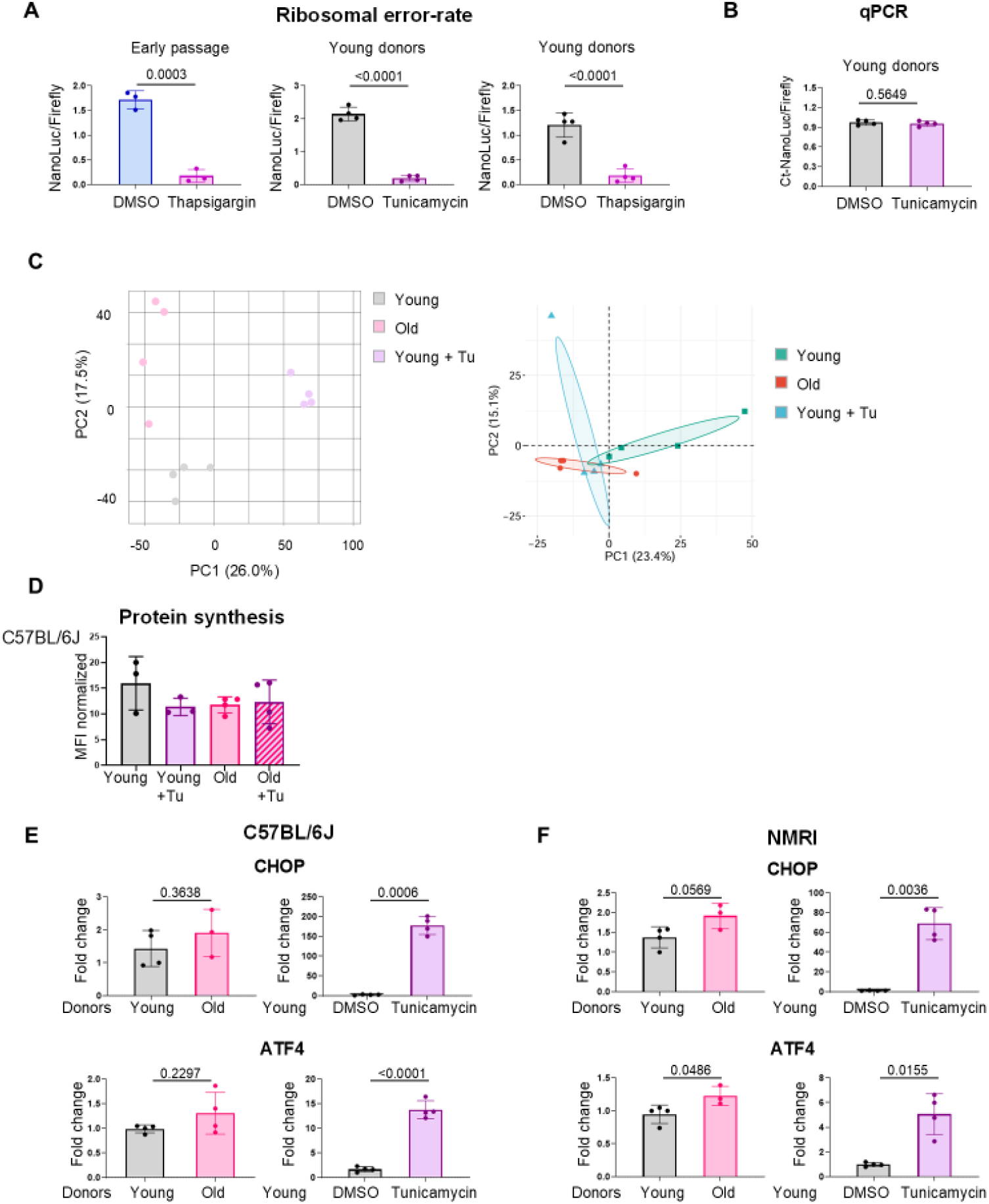
**A**. Translational fidelity assays, including treatment with thapsigargin (0.4 µM) and tunicamycin (1 µg/ml) in both models. N = 3, independent experiments, and n = 4, individual donors **B**. Similar expression levels of NanoLuc and Firefly Luciferase after transfection. N = 4, individual donors **C**. PCA analysis of the Nanopore sequencing data and the Mass Spectrometry data. **D**. Protein synthesis assay (OPP) of C57BL/6J mouse fibroblasts from young and old donors (n = 3), treated with tunicamycin (1 µg/ml for 24 hours). **E/F**. RT-qPCR analysis of C57BL/6J and NMRI mouse fibroblasts. Each data point represents an individual donor, n ≥ 3.

**figure S3.**
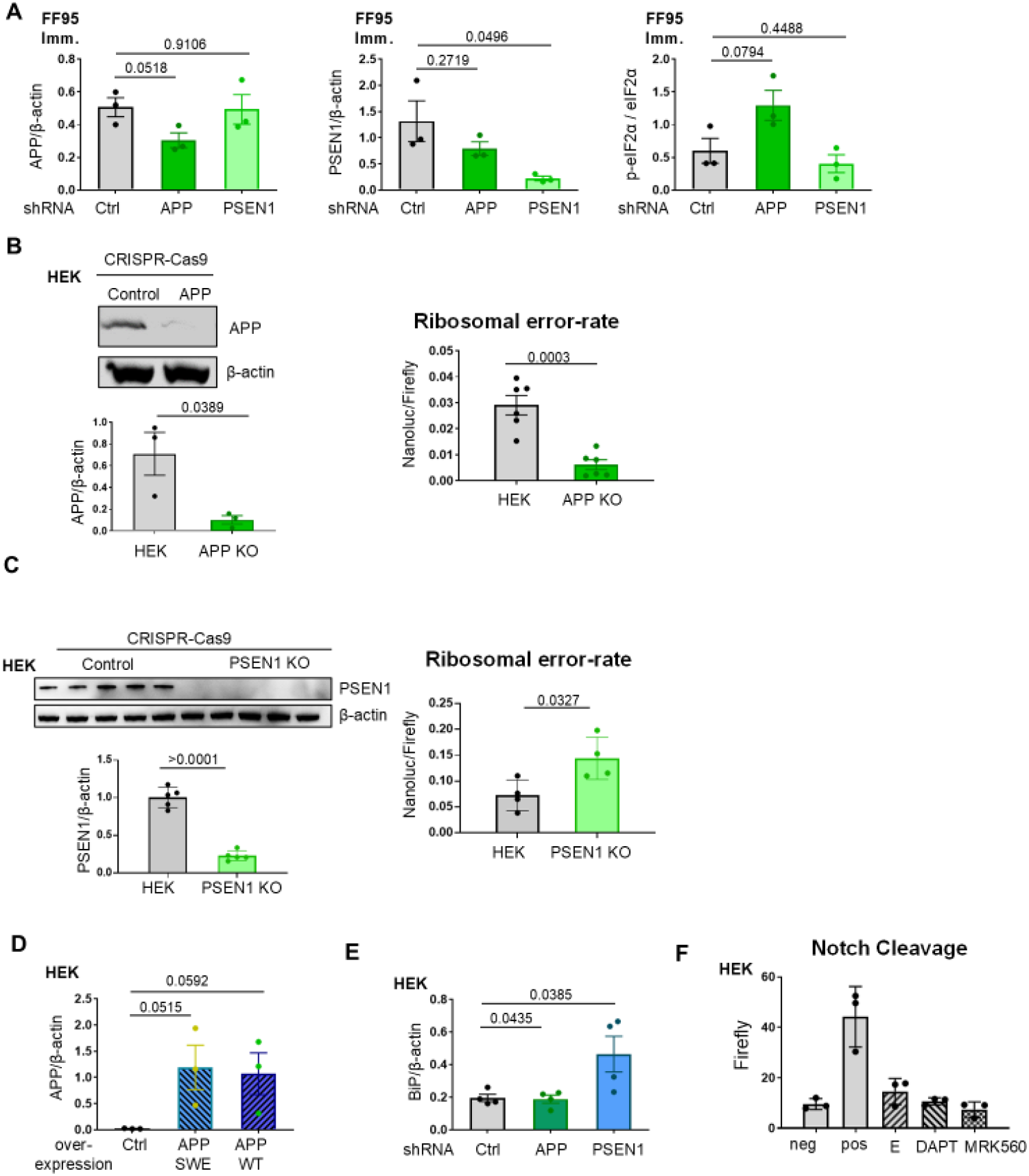
**A**. Western blot quantification (APP, PSEN1, peIF2α) of FF95 immortalized fibroblast extracts with shRNA expression targeting ctrl, APP, or PSEN1. N = 3, independent experiments. **B**. Verification of APP CRISPR-Cas9 knockout in HEK293T by Western blot (left). N = 3, independent experiments. Translational fidelity assay with stop codon readthrough reporter (right). N = 6, independent experiments. **C**. Verification of PSEN1 CRISPR-Cas9 knockout in HEK293T by Western blot (n = 5, independently cultured cells). Translational fidelity assay (right). N = 4, independent experiments. **D**. Western blot quantification of APP overexpression in HEK293T (n = 3, independent experiments). **E**. Western blot quantification of BiP protein level in HEK293T with stable expression of various shRNAs (n = 4, independent experiments). **F**. Verification of γ-secretase inhibitor activity (Compound E, DAPT, MRK560) using Notch cleavage assay (n = 3, independent experiments).

